# Bicistronic expression and differential localization of proteins in insect cells and *Drosophila suzukii* using picornaviral 2A peptides

**DOI:** 10.1101/766188

**Authors:** Jonas Schwirz, Ying Yan, Zdenek Franta, Marc F. Schetelig

**Affiliations:** Fraunhofer Institute for Molecular Biology and Applied Ecology IME, Winchesterstraße 2, Germany; Microscopy Core Facility, Institute of Molecular Biology gGmbH (IMB), Ackermannweg 4, Mainz, Germany; Justus-Liebig-University Gießen, Department for Insect Biotechnology in Plant Protection, Winchesterstraße 2, 35394 Gießen, Germany

**Keywords:** 2A peptides, localization signal, *Drosophila suzukii*, insect transgenesis, insect pest management

## Abstract

Polycistronic expression systems in insects can be used for applications such as recombinant protein production in cells, enhanced transgenesis methods, and the development of novel pest-control strategies based on the sterile insect technique (SIT). Here we tested the performance of four picornaviral 2A self-cleaving peptides (TaV-2A, DrosCV-2A, FMDV 2A1/31 and FMDV 2A1/32) for the co-expression and differential subcellular targeting of two fluorescent marker proteins in cell lines (*Anastrepha suspensa* AsE01 and *Drosophila melanogaster* S2 cells) and *in vivo* in the pest insect *Drosophila suzukii*. We found that all four 2A peptides showed comparable activity in cell lines, leading to the production of independent upstream and downstream proteins that were directed to the nucleus or membrane by a C-terminal nuclear localization signal (NLS) on the upstream protein and a poly-lysine/CAAX membrane anchor on the downstream protein. Two of the 2A peptides were inserted into *piggyBac* constructs to create transgenic *D. suzukii* strains, confirming efficient ribosomal skipping *in vivo*. Interestingly, we found that the EGFP-CAAX protein was distributed homogeneously in the membrane whereas the DsRed-CAAX protein formed clumps and aggregates that induced extensive membrane blebbing. Accordingly, only flies expressing the EGFP-CAAX protein could be bred to homozygosity whereas the DsRed-CAAX construct was lethal in the homozygous state. Our results therefore demonstrate that four different 2A constructs and two novel targeting motifs are functional in *D. suzukii*, and that DsRed-CAAX shows dosage-dependent lethality. These molecular elements could be further used to improve expression systems in insects and generate novel pest control strains.

**Highlights:**

- Four picornaviral 2A peptides have been studied for their self-cleaving ability in cell lines and *in vivo* in the pest insect *Drosophila suzukii*.
- All tested 2A peptides showed comparable activity that resulted in the production of independent upstream and downstream proteins.
- The proteins co-expressed by 2A peptides were either directed to the cell nucleus by a C-terminal nuclear localization signal (NLS), or to the cell membrane by a poly-lysine/CAAX membrane anchor.
- The combination of optimized membrane localization signals fused to DsRed generated an intrinsically lethal phenotype, which can be used to develop novel pest control strains.

## 1. Introduction

Spotted Wing Drosophila (*Drosophila suzukii*) is a devastating invasive pest that has spread throughout Asia, the Americas and Europe (http://www.cabi.org/isc/datasheet/109283), where it feeds and breeds on soft-skinned fruits such as strawberries, raspberries, elderberries and cherries [1–3]. The invasive success of *D. suzukii* partly reflects its life cycle. A single female can lay several hundred eggs, which she deposits directly into the fruit by penetrating the skin with a serrated ovipositor. Depending on the climate, eggs hatch within 72 h [2] and larvae feed inside the fruit as they develop through three instars until pupation. With more than 10 generations per year and the ability to multiply within a few weeks [4], *D. suzukii* can destroy entire fruit harvests in a matter of days under permissive environmental conditions. The short generation time and high fecundity of *D. suzukii* create challenging conditions for pest control, which has been a focus of scientific research for the last decade.

The sterile insect technique (SIT) is a pest control strategy that uses sterile males for population reduction. It works by releasing a large number of typically radiation-sterilized males into the environment, where they compete with unsterilized males to mate females. Mating between sterilized males and normal females produces unfertilized eggs, so the repeated release of sterile males causes population decline [5]. Only sterilized males contribute to the success of SIT, so the release of male-only populations makes SIT programs more efficient [6, 7]. However, the generation of genetic sexing strains is time consuming and labor intensive and must be established for every species from first principles. An alternative is the use of modern transgenic techniques to produce a transgenic embryonic sexing strain (TESS) based on the sex-specific splicing and early embryonic activation of pro-apoptotic genes [8–10]. The sex-dependent lethal effectors of sexing constructs can be adapted to various insect species, thus allowing the development of TESS strategies for many different pest insects in a reasonable timeframe. However, naturally-occurring mutations that restore viability would quickly spread when large numbers of individuals are produced and cannot be controlled by filter rearing systems [11] in a mass rearing facility, making additional pest control measures necessary [12]. Moreover, undetected resistance during the rearing process could lead to the establishment of a stable transgenic population in the field after release. Therefore, the robustness of TESS strategies must be ensured by creating built-in redundant systems that are independently lethal, which in turn requires a reliable strategy for the co-expression of multiple transgenes.

One straightforward approach that facilitates the co-expression of multiple proteins is the use of 2A self-cleaving peptides from Foot and mouth disease virus (FMDV) or other picornaviruses [13]. The 2A sequence is 18–22 amino acids in length and promotes ribosomal skipping such that the ribosome does not form the peptide bond between the glycine residue at the C-terminus of the 2A peptide and the N-terminal proline residue of the downstream protein [14, 15]. Because the skip occurs during translation, it allows the synthesis of both the upstream and downstream polypeptides, theoretically with an approximately equimolar ratio [16, 17]. Constructs containing 2A peptides have been used for biomedical research [18, 19], to achieve the production of multiple recombinant proteins in transgenic plants [20, 21], for the quantitative co-expression of marker proteins in fluorescence resonance energy transfer (FRET) applications [22], the improvement of CRISPR/Cas9 and TALEN genome editing methods [23, 24], and for the development of multigene expression systems in insects [25, 26].

Here, we developed a strategy to improve the current genetic control toolkit and to facilitate the production of multiple recombinant proteins in insects by testing the performance of four 2A peptides in insect cell culture systems (*Drosophila melanogaster* Schneider 2 (S2) cells and the *Anastrepha suspensa* cell line UFENY-AsE01) and in transgenic *D. suzukii* lines. As a model system, we co-expressed enhanced green fluorescent protein (EGFP) and the *Discosoma* red fluorescent protein (DsRed), with each protein carrying either a nuclear localization signal (NLS) from the *D. suzukii transformer* gene or a poly-lysine/CAAX motif for membrane localization [27, 28] at its C-terminus. We tested the efficiency of cleavage directed by the 2A peptides in insect cell cultures and discovered an unanticipated lethal effect caused by membrane-bound DsRed (DsRed-CAAX). This could be used as a universal lethal effector to replace pro-apoptotic genes, thus expanding the toolbox for transgenic pest control systems.

## 2. Materials and methods

### 2.1. Preparation of constructs

The NLS from the *D. suzukii transformer* (*Ds-tra*) gene was identified from transcriptomic data based on its similarity to the *D. melanogaster* NLS in the *pRed H-Stinger* construct [70, 71]. The NLS was amplified by PCR from vector pCR4-Ds-transformer_female_transcript using primers P260 and P226 (all primers are listed in Table S2). For DsRed-NLS, the *DsRed.T3* gene was amplified from #1425*_pBXL_attP220-PUbDsRedT3* [72] using primers P262 and P263. The amplified *DsRed* and *NLS* fragments were assembled using the Gibson Assembly Cloning Kit (NEB) following the manufacturer’s instructions. The product was re-amplified using primers P262 and P226, and transferred to the SacII and NotI restriction sites of the pIE4 overexpression vector (Novagen) to produce construct V125*_pIE4_DsRed-NLS*. For EGFP-NLS, the *EGFP* gene was amplified from #1419*_pBXL_PUbEGFP* [72] using primers P225 and P261. The amplified *EGFP* and *NLS* fragments were assembled by Gibson Assembly as above. The product was re-amplified using primers P225 and P226 and transferred to pIE4 as above, generating construct V126*_pIE4_EGFP-NLS*.

The CAAX motif was designed based on the Ras membrane anchor and contains a lysine-rich polybasic sequence followed by the C-terminal CAAX motif, where C = cysteine, A = any aliphatic amino acid, and X = any amino acid [27, 28]. *DsRed-CAAX* was amplified from #1425 using primers P264 and P265, the latter containing the CAAX motif. The amplified fragment was transferred to pIE4 as above to produce construct V123*_pIE4_DsRed-CAAX*. *EGFP-CAAX* was amplified from #1419 using primers P266 and P267, the latter containing the CAAX motif. The amplified fragment was transferred to pIE4 as above to produce construct V124*_pIE4-EGFP-CAAX*.

The TaV-2A, DrosCV-2A, Ta1/31-2A, and Ta1/32-2A sequences [16] were codon optimized for *D. suzukii* according to the Codon Usage Database (http://www.kazusa.or.jp/codon/) using Geneious Prime and corresponding oligonucleotides were transferred to cloning vector pEX by Eurofins Genomics. The cassettes were transferred to the SacII and NotI sites of vector V128*_pIE4-SV40* to generate the four 2A peptide destination vectors V142*_pIE4_DrosCV-2A_SV40*, V143*_pIE4_Ta1/31-2A_SV40*, V144*_pIE4_Ta1/32-2A_SV40*, and V145*_pIE4_TaV-2A_SV40*. *DsRed-NLS* was isolated from V125 and transferred to the SacII and AsiSI restriction sites of each 2A peptide destination vector, and *EGFP-CAAX* was then isolated from V124 and inserted at the ApaI and NotI sites of each 2A peptide destination vector containing *DsRed-NLS*, to generate the pIE4 constructs V150–V153 with *DsRed-NLS_2A_EGFP-CAAX* cassettes (Fig. 1B). To generate pIE4 constructs V154–V157 with *EGFP-NLS_2A_DsRed-CAAX* cassettes (Fig. 1B), the *EGFP-NLS* and *DsRed-CAAX* were isolated from V126 and V123, respectively, and inserted into V124 such that the localization signals were at identical positions but the EGFP and DsRed markers were exchanged. For the control construct V161*_pIE4_DsRed-NLS– EGFP-CAAX-SV40*, *EGFP-CAAX* was amplified from V124 using primers P362 and MFS8, and inserted into V125 at the AsiSI and NotI sites. For the control construct V162*_pIE4_EGFP-NLS-DsRed-CAAX-SV40*, *DsRed-CAAX* was amplified from V123*_pIE4_DsRed-CAAX* using primers P363 and MFS8, and inserted into V126*_pIE4_EGFP-NLS* at the AsiSI and NotI sites.

**Figure 1.**
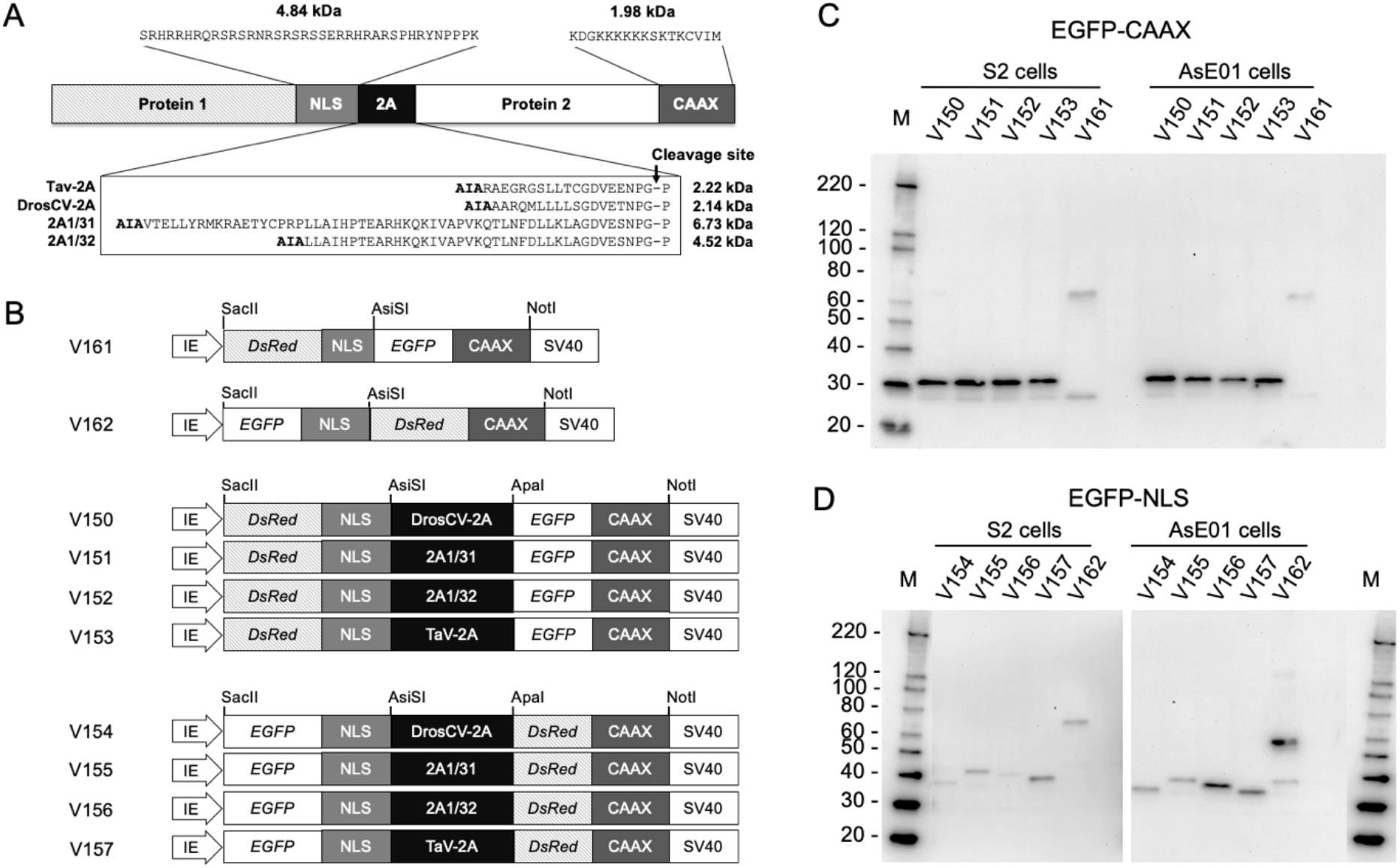
*In vitro* analysis of bicistronic constructs. A) The co-expression cassette contains one of four different 2A peptides, a nuclear localization signal (NLS) from the *D. suzukii transformer* gene, a polylysine-CAAX membrane tag (CAAX) and two proteins. The amino acid sequences and molecular weights of the NLS, CAAX, and the four different 2A peptides are indicated. B) Schematic drawing of pIE-4 derived overexpression vectors. Bicistronic constructs for EGFP-NLS (or DsRed-NLS) and DsRed-CAAX (or EGFP-CAAX), are either separated by a 2A peptide (V150–V157) or contain no 2A peptide (V161 and V162) and are expressed under the control of the immediate early IE promoter (IE). Unique restriction sites used for cloning are indicated. The AIA-sequences upstream of the 2A peptides are derived from an artificial AsiSI restriction site inserted for cloning purposes. Western blots are shown for EGFP-CAAX proteins (C) and EGFP-NLS (D). All constructs are cleaved correctly as demonstrated by the uniform size of CAAX-linked proteins (C), although NLS-tagged and cleaved proteins vary in size due to the variable length of the 2A cleavage site (D). The controls (V161 and V162) show a fusion protein of EGFP and DsRed of roughly double the size. M = marker (kD).

The *piggyBac* vector V204_*pBXL-PUbAmCyan_PUbDsRed-NLS-SV40* was then prepared by cloning a part of the 5′ region of *DsRed* together with the SV40 region from V193_*pIE4-Ds-traIntron-DsRed-NLS-SV40* into V198_*pBXL-attP-[PUbAmCyan_rev]_PUbDsRed-NLS* at the SbfI and BglII sites and then flipping the *PUb-AmCyan-SV40* cassette by cutting the vector with Bsp119I and re-ligating the insert. Construct V198 was prepared by transferring the *DsRed_NLS* sequence from V125 to the SacII and BglII sites of V195_*pBXL-attP-PUbAmCyan_PUb_MCS* using primers P395 and P396. The co-expression cassettes were amplified from constructs V150, V153, V154, and V157 by PCR using primers P227 and P191, and the products were inserted to the SacII and BglII sites of V204, respectively, to obtain the final *piggyBac* transformation vectors V220–V223 (Fig. 3A).

### 2.2. Cell culture experiments

The *A. suspensa* cell line UFENY-AsE01 (AsE01) [73] and *D. melanogaster* S2 cells were grown in Schneider’s medium with 10% fetal bovine serum (FBS) and 1% penicillin/streptomycin in closed capped flasks without CO_2_ at 27.5°C and 25°C, respectively. For transfection experiments, the Xfectin transfection reagent (Takara) was used according to the manufacturer’s instructions. We seeded 4 × 10^5^ cells (live cell count) into 35-mm glass-bottomed cell culture µ-dishes (ibidi) in 400 µl of medium. After allowing 3 h to settle, the medium was removed and the cells were transfected for 4 h by adding 2 µg of plasmid DNA, 0.6 µl Xfectin and 37.4 µl Xfectin buffer made up to 400 µl in serum-free Schneider’s medium. The transfection medium was then replaced with 2 ml fresh standard growth medium (Schneider’s with 10% FBS and 1% penicillin/streptomycin). The AsE01 and S2 cells were incubated for approximately 14 h at 27.5°C and 25°C, respectively, then either fixed for 15 min in 4% paraformaldehyde in PBS for microscopy or harvested for total cell protein isolation as described below.

### 2.3. Imaging

Cells and tissues were imaged using an SP5 inverted confocal microscope (Leica Microsystems) equipped with 488 and 561 nm laser lines for EGFP and DsRed excitation, respectively. Acquired images were deconvolved using Huygens Essential v15.05 (Scientific Volume Imaging). Transient expression in G_0_ larvae and stable expression in transgenic heterozygous larvae was monitored using a M205FA microscope (Leica Microsystems) with filter sets CFP for AmCyan (ex. 436/20; em. 480/40), YFP for EGFP (ex. 510/20; em. 560/40), TxRed for DsRed (ex. 545/30; em. 620/60) and GFP-LP for overlay (ex. 425/60; em. 480). All settings (e.g. exposure time, gain, magnification) were identical for each filter so that the fluorescence intensity could be visually compared across images.

### 2.4. Insect rearing and germ-line transformation

Wild-type (USA strain) and transgenic *D. suzukii* lines were maintained at 25°C and 60% humidity with a 12-h photoperiod. Flies were anesthetized with CO_2_ for screening and to set up crosses. For germ-line transformation, eggs were collected over a 30-min period on grape juice agar plates (1% agar, water:grape juice ratio 7:3), desiccated for 10 min and then overlaid with halocarbon oil 700 (Sigma-Aldrich) as previously described [74]. A mixture of the *piggyBac* donor construct (500 ng/µl) and either the *phsp-pBac* transposase (200 ng/µl), or *phsp-pBac* and hyperactive transposase [75] mix (200 ng/µl each), or synthetic *piggyBac* RNA (200 ng/µl) helper plasmid (Table S1) was injected into wild-type embryos. Hatched G_0_ larvae were screened for transient AmCyan expression by epifluorescence microscopy. Surviving G_0_ adults were then backcrossed to wild-type males or virgin females with putative G_1_ transformant progeny selected by AmCyan fluorescence. Independent homozygous strains were established by single-pair inbreeding for successive generations and segregation test analysis of transformants outcrossed to wild-type flies.

### 2.5. Quantitative real-time PCR

Total RNA was isolated from heterozygous third-instar larvae using the ZR Tissue and Insect RNA MicroPrep kit (Zymo Research) and 1 μg of RNA was transcribed into cDNA using the iScript cDNA Synthesis Kit (Bio-Rad). Primer pairs MFS17/MFS18, P16/P17 and MFS19/MFS20 were designed to compare the expression levels of *EGFP* and *DsRed* to the *AmCyan* control. The iQ SYBR Green Supermix (Bio-Rad) was used for qPCR with 100 ng cDNA in a CFX96 Touch Real-Time time PCR Detection System (Bio-Rad). All reactions were performed on three biological and technical replicates. AmCyan was used as reference gene, and the 2^−∆∆Ct^ method [76] was used with all samples relative to the DsRed expression in V220. One-way ANOVA (SigmaPlot v12.5) was used to compare the relative expression levels of target genes and means were separated using Duncan’s method.

### 2.6. Total protein extraction

AsE01 and S2 cells were grown and transfected as described above. After 14 h of incubation, cells were harvested in sterile PBS and centrifuged (30 min, 2716 rcf, 10°C for AsE01; 10 min, 244 rcf 10°C for S2). Cell pellets were washed with 1 ml ice-cold PBS and centrifuged again as above. The pelleted cells were lysed in ice-cold Pierce RIPA buffer (Thermo Fisher Scientific) supplemented with 1x SIGMAFAST Protease Inhibitor (Sigma-Aldrich). The cell mixture was vigorously pipetted, supplemented with 4x Laemmli sample buffer (Bio-Rad) containing 50 mM DTT and boiled for 5 min at 95°C. For transgenic flies, 15 larvae (heterozygous third instar) from each line were homogenized in 2x Laemmli sample buffer (Bio-Rad) containing 355 mM 2-mercaptoethanol (Sigma-Aldrich). Homogenates were incubated at 95°C for 5 min then centrifuged at 10.000 g for 5 min. The supernatant was stored at –20°C.

### 2.7. SDS-PAGE and western blot analysis

Cell lysates were resolved by reducing SDS-PAGE and stained with colloidal Coomassie Brilliant Blue (Carl Roth) or transferred to a polyvinylidene difluoride (PVDF) membrane (Merck Millipore) using the TransBlot Turbo Transfer System (Bio-Rad). The membrane was blocked overnight in Tris-buffered saline containing 0.1% Tween-20 (TBST) and 2.5% nonfat dried milk (Sigma-Aldrich) and washed (3x 10 min) in TBST before incubating at room temperature for 2 h with polyclonal or monoclonal antibodies against EGFP (AB290 from ABCAM) or DsRed (AB62341 and AB28664 from ABCAM for cell culture samples, and M204-7 from Medical & Biological Laboratories for fly tissues) diluted 1:2000 in phosphate-buffered saline with 0.05% Tween-20 (PBST) containing 2.5% nonfat dried milk. The membrane was washed with TBST (3x 10 min) and incubated with the secondary goat anti-rabbit whole IgG horseradish peroxidase (HRP) conjugated antibody (Dianova) diluted 1:5000 in TBST containing 2.5% nonfat dried milk) for 1 h at room temperature. The signal was developed using Lumi-Light Plus Western Blotting Substrate (Roche) and visualized using a VersaDoc Imaging System (Bio-Rad).

## 3. Results

### 3.1. Construct design

We designed a range of bicistronic expression constructs for *in vitro* cell culture and *in vivo* expression in transgenic *D. suzukii* lines incorporating 2A self-cleaving peptides derived from two insect viruses, namely *Thosea asigna* virus (TaV) and *Drosophila* C virus (DrosCV), as well as two artificial 2A sequences derived from FMDV (2A1/31 and 2A1/32) [16] (Fig. 1A,B). The amino acid sequences were identical to those used previously [16] but the nucleotide sequences were codon optimized for *D. suzukii*. Each test construct contained an upstream *DsRed* and downstream *EGFP* gene (or vice versa) separated by one of the 2A peptides, which increased the size of the upstream protein by 2.22–6.73 kDa depending on the precise cleavage site. The upstream gene in each construct carried a C-terminal NLS from the *D. suzukii transformer* gene whereas the downstream gene carried a C-terminal poly-lysine/CAAX membrane tag to avoid potential interference with the 2A sequence [27]. These tags increased the size of the proteins by another 4.84 kDa (NLS) and 1.98 kDa (poly-lysine/CAAX), respectively. The attachment of different tags to the upstream and downstream proteins in each construct directed the two proteins to different subcellular sites. Control constructs were designed as stated above but lacking a 2A sequence, thus generating conventional DsRed-EGFP and EGFP-DsRed fusion proteins. Expression in S2 cells was driven by the immediate early promoter of the pIE-4 vector. Each of the four 2A peptides was tested in combination with DsRed-NLS + EGFP-CAAX (Fig. 1A, V150–V153) and EGFP-NLS + DsRed-CAAX (Fig. 1A, V154–V157), leading to eight experimental construct designs plus two fusion protein controls (Fig. 1A, V161–V162).

### 3.2. The 2A peptides achieve efficient multigene expression in cell lines

The protein cleavage activity for each construct in cell culture was determined by western blot (detection of EGFP or DsRed) and fluorescence microscopy. Western blot analysis (Fig. 1C) revealed single bands corresponding to the anticipated size of EGFP for each of the bicistronic constructs, whereas bands of approximately double the size were detected for the control construct, corresponding to the size of the EGFP-DsRed fusion protein. We observed slight size variations when comparing constructs V154, V155, V156 and V157 (EGFP-NLS) to V150, V151, V152 and V153 (EGFP-CAAX) reflecting the size difference between the NLS (4.84 kDa) and CAAX tag (1.98 kDa) (Fig. 1C, D), as well as smaller differences within these sets of constructs reflecting the lengths of the four different 2A peptides (Fig. 1D). This was made possible by using the sensitive and specific EGFP antibody. However, two different DsRed antibodies (AB62341 and AB28664) using different staining conditions failed to identify any specific DsRed products (Fig. S1).

Fluorescence microscopy confirmed that all constructs containing a 2A peptide achieved the distinct localization of the nuclear and membrane-targeted proteins in AsE01 cells (Fig. 2) and S2 cells (Fig. S2). This indicates the discrete and consecutive production of both proteins during translation and hence functional ribosomal skipping. In contrast, EGFP and DsRed were co-localized in cells transfected with the control constructs V161 and V162, although we also detected regions of green fluorescence without (or with negligible) red fluorescence, perhaps due to the longer maturation time of the DsRed.T3 protein compared to EGFP, and the initial green fluorescence of immature DsRed protein [29]. The control construct fusion proteins were not localized to the inner nucleus but were present in the plasma membrane and nuclear membrane, probably due to competition between nuclear import and membrane anchoring by the CAAX tag. For the construct with DsRed-NLS and DrosCV-2A (V150), we observed nuclei with spots of strong DsRed fluorescence in most cells, potentially representing subnuclear compartments, contrasting with the homogenous nuclear distribution observed for the other 2A peptides. The cells expressing construct V150 were shrunken and showed evidence of membrane blebbing in the EGFP channel, which may indicate the onset of apoptosis caused by cytotoxic DsRed accumulation in the nucleus. We did not observe such spots in cells expressing construct V154 (EGFP-NLS_DrosCV2A_DsRed-CAAX), suggesting that the effect is unique to the combination of DsRed-NLS and DrosCV-2A. For constructs V154 and V157, EGFP was not entirely restricted to the nucleus and a considerable amount was also found in the membrane, albeit not precisely co-localized with DsRed. Furthermore, in cells transfected with these constructs, EGFP expression was still strong in the nucleus, whereas no EGFP was found in the nucleus for the control construct (V162). These results indicated that both fluorophores were expressed independently with all four 2A peptides we tested.

**Figure 2.**
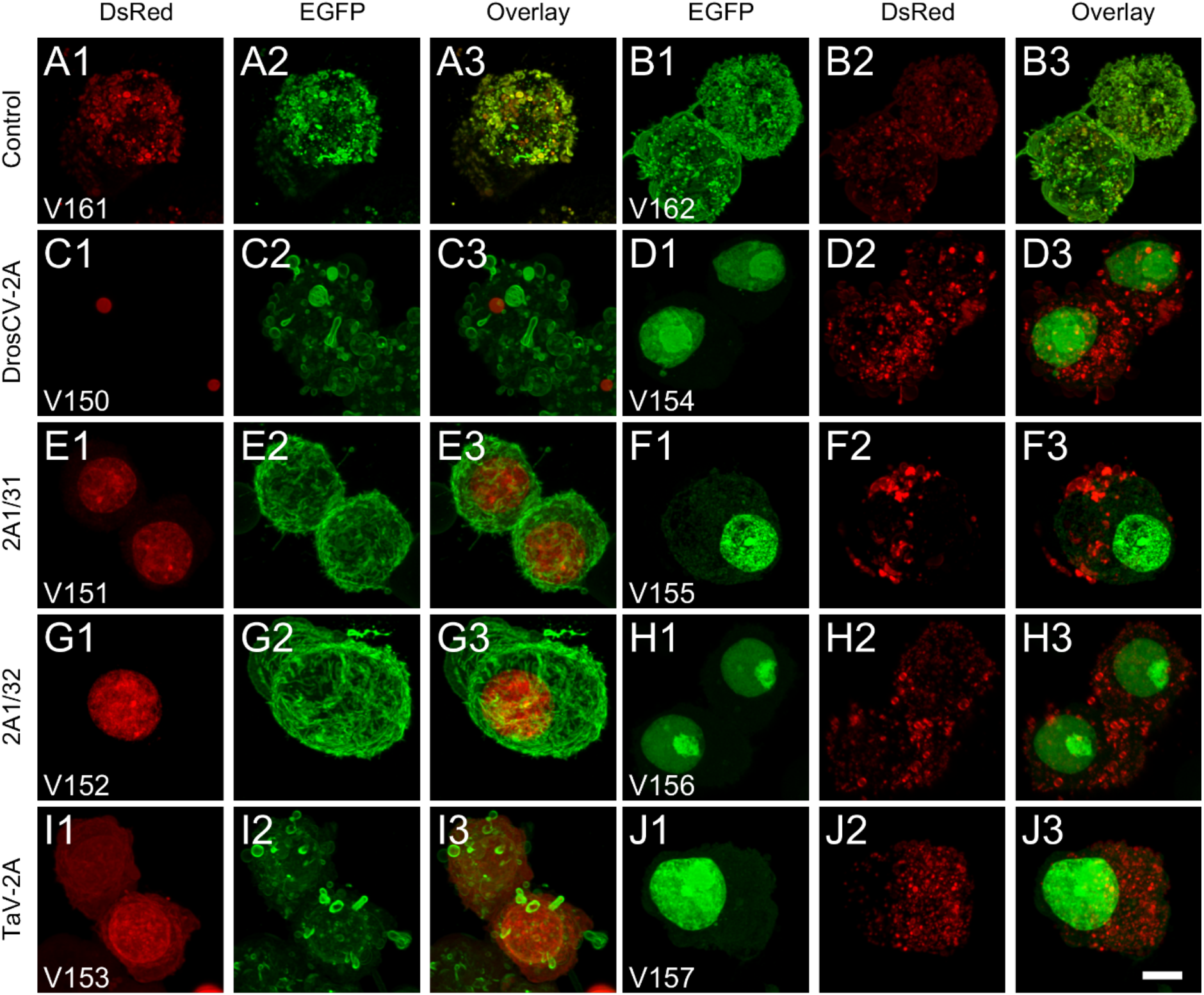
Protein expression using 2A peptides in AsE01 Tephritid cells. The control constructs V161 and V162 (A1-A3 and B1-B3) expressed an EGFP-DsRed fusion protein and the two fluorophores co-localized in membrane regions. Weaker or undetectable DsRed in some areas may be due to the longer maturation time of DsRed. In the constructs containing a 2A peptide (C1-J3), EGFP and DsRed were independently localized due to ribosomal skipping during the translation process. The EGFP-CAAX protein (C2, E2, G2, and I2) was usually homogenously localized to the cell membrane, whereas DsRed-CAAX (D2, F2, H2 and J2) in contrast formed clusters. In the construct containing the DrosCV-2A peptide (V150, C1-C3), the nuclear DsRed mostly showed subnuclear agglomerations of strong fluorescence instead of DsRed distribution throughout the nuclear region. These cells often showed a membrane blebbing effect in the EGFP channel. A similar, but weaker phenotype was observed for the V150 and V153 constructs (C1-C3, I1-I3). Scale bar = 5 µm.

### 3.3. DsRed-CAAX may trigger the loss of membrane integrity

Whereas EGFP-CAAX was always homogenously distributed in the cell membrane, forming a sheath that decorated the cell border, DsRed-CAAX formed unexpected clusters observed as intense fluorescent spots in the cell membrane, interspersed with DsRed-free regions. This was also true for the control constructs V161 and V162, which contained a C-terminal membrane tag, but due to the direct fusion of EGFP and DsRed in these constructs we also observed discrete spots of EGFP fluorescence (Fig. 2). The basis for this clustering is unclear, but may reflect the self-association of DsRed to form obligate tetramers *in vitro* and *in vivo* [30]. The cells expressing membrane-bound DsRed often showed extensive membrane blebbing, suggesting that the DsRed aggregates were detrimental, as previously reported in bacteria [31]. Similarly, we found that *Escherichia coli* transformed with the DsRed-CAAX constructs grew slowly and produced low quantities of DNA, supporting the hypothesis that DsRed-CAAX inhibits cell growth and survival.

### 3.4. The 2A peptide sequences are also active in vivo in D. suzukii

Four *piggyBac* constructs (V220–V223) containing different co-expression cassettes (Fig. 3A) were injected into *D. suzukii* embryos at the early blastoderm stage. The transient expression of EGFP and DsRed (mediated by either DrosCV-2A or TaV-2A) was detected during late embryogenesis and continued to the third-instar stage (Figure 3B), showing that the 2A peptides are compatible with the DmPUb promoter for fast and efficient transient expression. Using *piggyBac*-mediated transformation, we generated independent transgenic lines expressing constructs V220 (line V220), V221 (line V221) and V223 (lines V223-M2 and V223-F2) (Table S1), all of which showed constitutive expression of AmCyan, EGFP and DsRed (Fig. 3B). The intensity of AmCyan fluorescence differed among the transgenic lines, possibly due to genomic position effects. The intensity of EGFP and DsRed fluorescence also differed among the transgenic lines, probably reflecting the combined impact of genomic position effects and position within the construct (before or after a 2A peptide). The V220 and V221 lines were bred to homozygosity and were stably maintained for more than one year. In contrast, the V223 lines (in which TaV-2A was used for DsRed-CAAX expression) were homozygous lethal and could only be maintained as heterozygous populations. Multiple attempts to generate V222 transgenic flies using different transposase helpers failed (Table S1), possibly due to the extensive membrane blebbing triggered by the DrosCV-2A and DsRed-CAAX combination, as observed in the cell cultures. All transgenic lines showed localization and expression of EGFP and DsRed in tissues such as gut, Malpighian tubules and accessory glands. The cardia (junction between the foregut and midgut) was prepared for confocal imaging as a representative example because it has a large surface and is easy to handle. Cells from lines V220 and V221 were intact and showed clear and homogenous nuclear DsRed-NLS localization as well as a membrane-bound EGFP-CAAX signal. But in line V223, EGFP-NLS and DsRed-CAAX fluorescence were distributed heterogeneously across the cardia and formed clusters, accompanied by cell shrinkage and membrane blebbing (Fig. 4). These results suggest that DsRed-CAAX expression is detrimental to flies, and the effect is more profound when driven by DrosCV-2A rather than TaV-2A.

**Figure 3.**
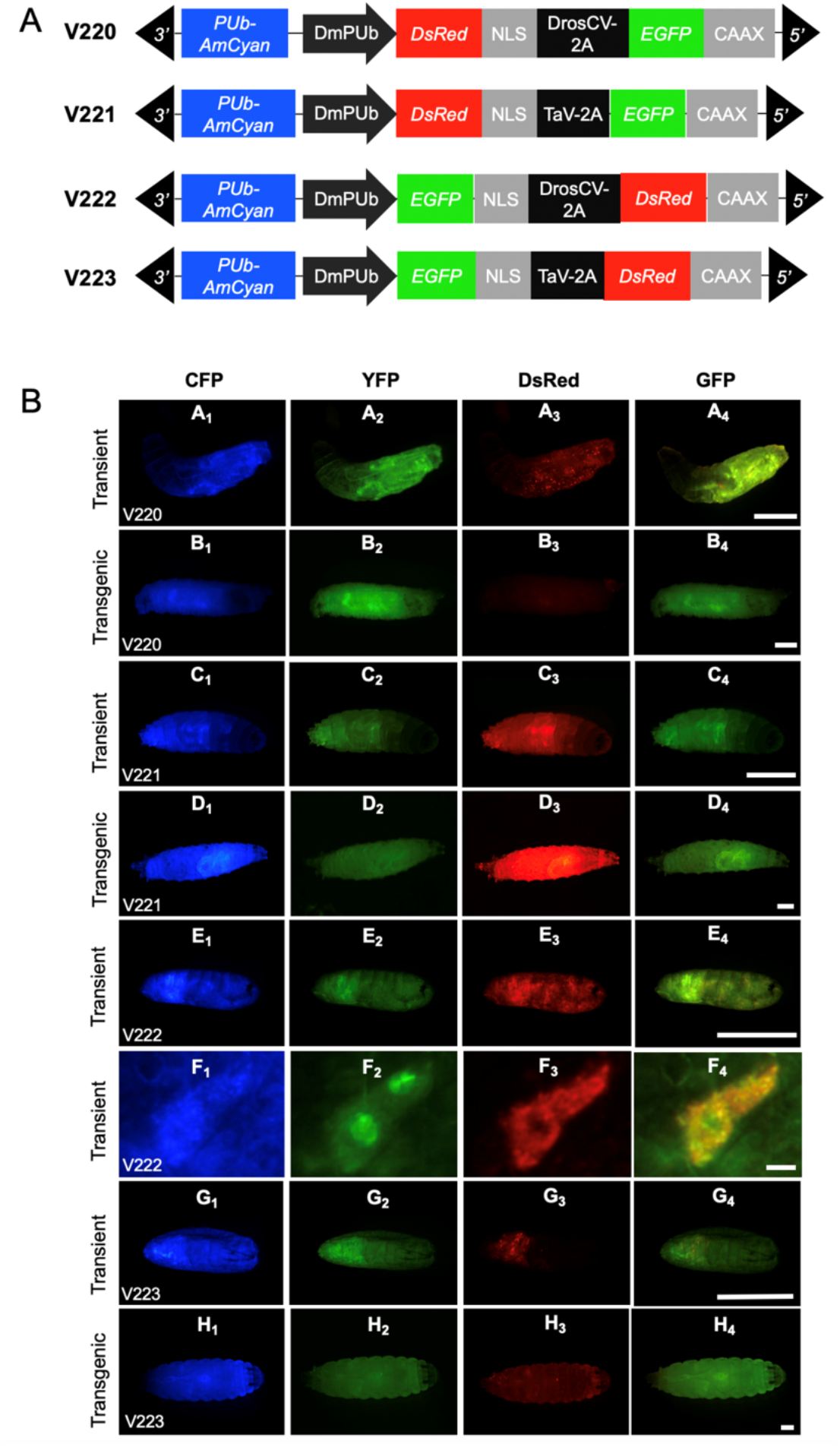
Transient and stable expression of fluorescent proteins mediated by 2A peptides in *Drosophila suzukii*. (A) Schematic drawing of vectors used for germline transformation (the size of the boxes is not to scale). Bicistronic constructs for EGFP-NLS (or DsRed-NLS) and DsRed-CAAX (or EGFP-CAAX), separated by a 2A peptide, expressed under the control of the *D. melanogaster* PUb promoter (DmPUb). The constructs also contain an AmCyan gene under the control of the DmPUb promoter as a transformation marker. (B) The fluorescent markers under the control of the DmPUb promoter are constitutively expressed in larvae following transient expression (A, C, E, F, G) and in stable transgenic lines (B, D, H). Both DrosCV-2A and TaV-2A efficiently co-expressed EGFP and DsRed in transient expression experiments (G_0_, the first/second instar larvae) and in transgenic flies (heterozygous, the third instar larvae). We were unable to recover homozygous transgenic lines expressing V222, but clear localization of EGFP and DsRed was observed by transient expression (F, gut tissue; scale bar = 10 µm). All other scale bars = 0.5 mm. The intensity of AmCyan, EGFP and DsRed fluorescence differs among transgenic lines, possibly due to genomic position effects and positioning of the markers within the construct.

**Figure 4.**
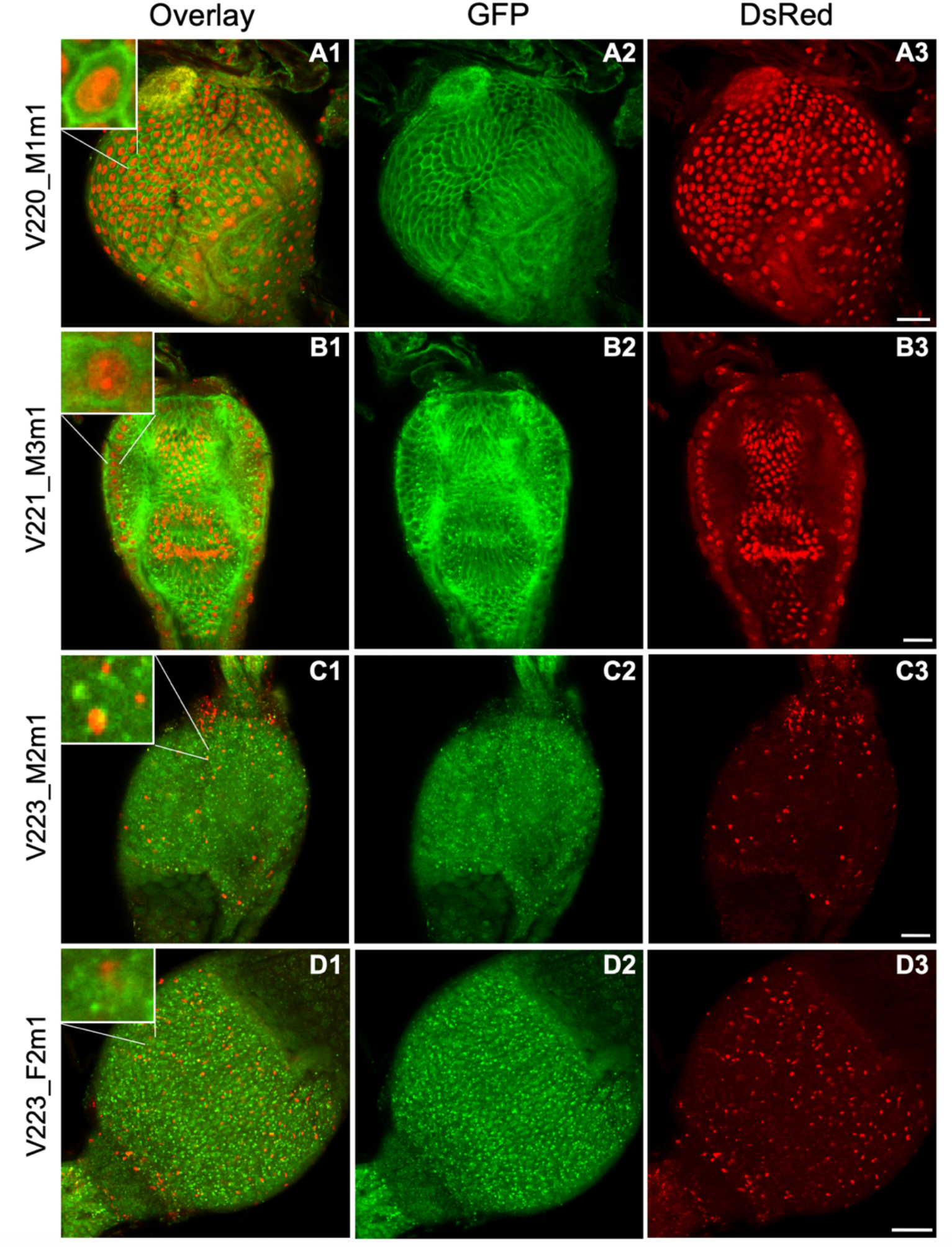
Stable expression of localized fluorescent proteins mediated by 2A peptides in transgenic *Drosophila suzukii*. Each cardia was dissected from third-instar larvae (heterozygous for the transgene) and used for imaging. Both EGFP and DsRed expression can be observed in the cardia from all four transgenic lines. V220_M1m1 and V221_M3m1 tissue shows membrane-localized EGFP-CAAX (A2, B2) and nuclear DsRed-NLS (A3, B3) expression. Lines V223_M2m1 and V223_F2m1 containing EGFP-NLS (C2, D2) and DsRed-CAAX (C3, D3) showed scattered clusters of fluorescence distributed across cells. Insets show individual cells. Scale bars = 20 µm.

### 3.5. Gene expression in constructs containing different 2A sequences

Finally, we carried out quantitative real-time PCR (qPCR) experiments to compare relative mRNA levels in transgenic *D. suzukii* lines carrying different 2A peptides and localization signals (Fig. 5A). EGFP-CAAX was expressed at 3.5-fold and 3.4-fold higher levels than DsRed-NLS in lines V220 (*P<0.001*) and V221 (*P<0.001*), respectively. Furthermore, DsRed-CAAX was expressed at 4.5-fold and 4.2-fold higher levels than EGFP-NLS in lines V223-M2 (*P=0.006*) and V223-F2 (*P=0.004*), respectively. Given that the cassette was, in each case, regulated by a single promoter (DmPUb) and the ribosomal skip takes place during translation, the significant difference between EGFP and DsRed mRNA levels in the same transgenic line is more likely to be associated with the localization signals. In general, the gene upstream of the 2A peptide produces more protein than the downstream gene due to the molar excess of the N-terminal cleavage product [16, 17]. Here, mRNA levels of EGFP-CAAX (downstream of the 2A peptide) in lines V220 and V221 were 9.8-fold (*P<0.001*) and 9.1-fold (*P<0.001*) higher than those of EGFP-NLS (upstream of the 2A peptide) in line V223-M2, respectively, and 8.6-fold (*P<0.001*) and 8.0-fold (*P<0.001*) higher than those of EGFP-NLS in line V223-F2, respectively (Fig. 5A). Accordingly, the EGFP-CAAX protein produced uniform-sized bands in lines V220 and V221, but EGFP-NLS was not detected in lines V223-M2 and V223-F2 (Fig. 5B). Thus, the localization signal may play a more important role than the position (upstream or downstream of the 2A peptide) in determining the mRNA and protein levels for EGFP. In addition, EGFP-CAAX was cleaved more efficiently in the line containing DrosCV-2A (V220) rather than TaV-2A (V221) based on the intensity of the protein signal (Fig. 5B), which is consistent with the germ-line transformation results of V222 (DrosCV-2A/DsRed-CAAX) and V223 (TaV-2A/DsRed-CAAX). On the other hand, there was no significant difference in DsRed mRNA levels between lines V220/V221 and lines V223-M2/V223-F2 (*P=0.223* for V220 vs V223-M2, *P=0.140* for V220 vs V223-F2, *P=0.198* for V221 vs V223-M2, *P=0.119* for V221 vs V223-F2; Fig. 5A). However, the DsRed-CAAX protein (downstream of the 2A peptide) in line V223 was uniformly cleaved, as indicated by the single band, whereas DsRed-NLS (upstream of the 2A peptide) was not detected in lines V220 and V221 (Fig. 5B). A recent study showed that position does not affect EGFP and DsRed mRNA levels, but translation was enhanced for DsRed alone when placed downstream of the 2A peptide [32]. Both the localization signal and the position relative to the 2A peptide therefore appear to influence the level of DsRed protein.

**Figure 5.**
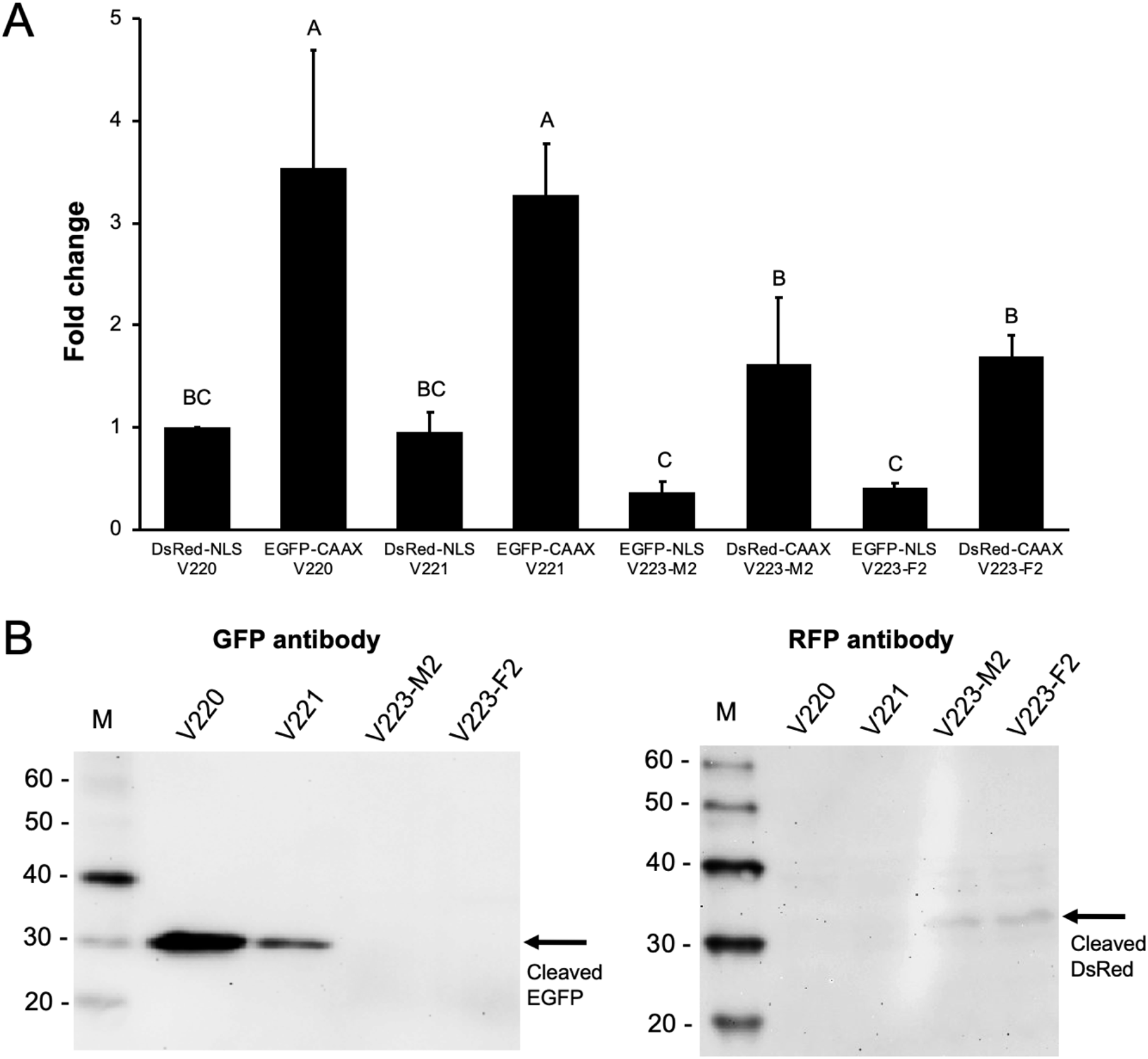
Levels of mRNA and protein in bicistronic transgenic *Drosophila suzukii* adults. (A) Fusion gene expression (mRNA level) determined by real-time PCR. The levels of EGFP and DsRed mRNA are relative to AmCyan mRNA as an internal control for each transgenic line. Data are means ± SE (n = 3) and bars with different letters are significantly different at p<0.05 (one-way ANOVA, Duncan’s method). (B) Analysis of EGFP and DsRed protein levels by western blot. Protein was extracted from the whole body of third-instar larvae (n = 15). The CAAX-tagged EGFP (downstream of the 2A peptide) from homozygous V220 and V221 flies was detected and cleaved to a uniform size, as indicated by the single band detected with the anti-GFP antibody (AB290). The NSL-tagged EGFP (upstream of the 2A peptide) from heterozygous V223 flies (the homozygous state was lethal) was not detected under the same conditions. The CAAX-tagged DsRed (downstream of the 2A peptide) from V223 flies was detected and cleaved to a uniform size, as indicated by the single band detected with the anti-RFP antibody (M204-7). The NLS-tagged DsRed (upstream of the 2A peptide) from lines V220 and V221 was not detected under the same conditions. M = marker (kD).

## 4. Discussion

In picornaviruses, 2A peptide sequences enable the production of multiple virus proteins from a single, polycistronic mRNA [13]. The sequence causes the ribosome to pause at its C-terminal glycine residue, followed by the re-initiation of protein synthesis at the subsequent proline residue, leading to the formation of separate upstream and downstream proteins [14, 15]. We tested four 2A peptides: two natural peptides from insect viruses (TaV-2A and DrosCV-2A) and two synthetic peptides based on the natural peptide in FMDV (2A1/31 and 2A1/32) [16], for co-expression of two marker gene EGFP and DsRed. According to quantitative expression data, the mRNA sequences of the markers are significantly different expressed depending on their position and/or localization tag in the multicistronic mRNA. This is an unexpected finding, because the measurement at two loci of a bicistronic mRNA should result in the same amount of transcript. In the presented measurements, the downstream cassette carrying the CAAX membrane tag always showed a stronger signal. This phenomenon was analysed under two hypotheses. First, the downstream part genuinely is more abundant, which could be caused by the upstream part degrading faster than the downstream part leaving more of the template. In that respect, the overall RNA quality of all samples was checked and was consistent. Also, the primer binding sites are towards the 3’ end of each marker gene. So, this would require a degradation of almost the complete first cistronic cassette. While this is rated as unlikely, the possibility still exists. Second, the two parts are equally abundant, as would be expected given they are meant to be part of one molecule, but the signals are not comparable for some other currently unexplainable reasons. For that, amplification efficiencies during qPCR were in a range of 0.9 to 1.1 for all samples and positions on the transcript and could therefore not be a technical error. Another option could be the localization signal sequences, if those sequences could influence amplification levels during qPCR. Fluorescent intensities (Fig. 4) speak against such a hypothesis and suggest an equal or more abundant expression of the upstream cassettes. Taken together, currently the phenomenon cannot be explained with certainty.

On a protein level, all four peptides showed comparable activity in *A. suspensa* AsE01 and *D. melanogaster* S2 cells, leading to the production of independent upstream and downstream proteins that were directed to the nucleus and membrane respectively, due to the addition of a C-terminal NLS to the upstream protein and a poly-lysine/CAAX motif to the downstream protein. The synthesis of independent proteins was confirmed by western blot (revealing bands of the anticipated size for the separate EGFP and DsRed polypeptides) and immunofluorescence analysis of cells and tissues (revealing distinct subcellular locations for each fluorophore). Several other 2A peptides could be potential candidates for future studies in *D. suzukii*, including the Porcine teschovirus-1 2A peptide, which has been used in *D. melanogaster* [25].

The use of 2A peptides brings significant advantages that are beneficial for pest control applications. First, because the initiation of translation is the rate-limiting step in protein synthesis [33], polycistronic constructs in which genes are separated by 2A peptides increase the overall efficiency of translation by yielding more proteins per initiation event [16, 25]. Such robust expression can increase the efficiency of functional constructs that are employed in a pest control strategy. Second, translation using a multigene construct can achieve a certain ratio of the proteins related to the positions of the 2A peptide [17, 32], and is therefore ideal for the co-expression of gene products that work cooperatively and require an optimal stoichiometric ratio, such as the pro-apoptotic genes used for TESS [34, 35]. Liu et al demonstrated that the position and combination of proteins can alter the expression strength of proteins in cells [17]. So, the exact ratio for each combination and tissue needs to be evaluated. Third, the small size of 2A peptides means that they have little impact on transgene size. This is a significant advantage in complex multigene constructs where the size of multiple promoters or internal ribosome entry sites (IRES) can increase the overall insert size by several kilobase pairs, which reduces the efficiency of bacterial transformation and transposon/recombination-mediated transformation in insects [36–39]. Specifically, the small sizes of 2A peptides make them ideal for the development of safer and more effective SIT approaches that include redundant lethal effectors to prevent the restoration of fertility or viability by mutation. For example, some pro-apoptotic genes are rather small such as *head involution defective* (*hid*) at 732 bp and *reaper* (*rpr*) at 380 bp [34] and have been tested successfully in agricultural pest insects to induce male sterility [40], embryonic lethality [41–43] and TESS [8–10]. Similarly, multiple copies of effector molecules such as the *tetracycline transactivator* (*tTA*) in the releasing insects that carry dominant lethals (RIDL) system [44, 45] or sex determination gene in the sex-ratio distortion (SRD) system [46] could be co-expressed by 2A peptides in pest insects. With several copies of an effector gene on one construct, the desired effect could be enhanced, and the system could cope with the loss of individual genes due to missense mutations, ensuring that the emergence of resistance was delayed. However, 2A peptide constructs need an intact, continuous open reading frame and are therefore susceptible to nonsense or frameshift mutations affecting the upstream genes. An optimal backup system would therefore consist of two or more independent multigene constructs with individual genes separated by 2A peptides.

Our investigation of the performance of 2A peptides also led to the development of new tools that could be useful in transgenic *D. suzukii* systems. One such tool was the *Ds-tra* NLS, which successfully directed tagged proteins to the nucleus and could be used, for example, to shuttle effector proteins encoded by early embryonic lethality constructs, or to increase the efficiency of transgenesis and genome editing by directing the corresponding enzymes to the nucleus. The presence of a NLS was shown to improve genome editing efficiency when placed at the N-terminus or C-terminus (or both) of Cas9 [47]. Another useful tool was the CAAX tag, which anchored the fluorescent proteins to the plasma membrane or to the membranes of subcellular compartments. Remarkably, when this tag was combined with DsRed.T3, the membrane-bound protein formed aggregates that appeared to have a detrimental effect on cells both *in vitro* and *in vivo*. Cytotoxicity has been reported and studied for several DsRed variants before [29, 48] and tagged versions of DsRed.T3 showed toxicity in *E. coli* cells [49]. The mechanism for toxicity for DsRed variants is not entirely understood [50] and might vary between cell types or species [51]. Our cell culture experiments showed a detrimental effect when overexpressing the DsRed.T3 with the membrane tag CAAX. While this is in line with other reports of DsRed toxicity, we have not seen such strong effects of DsRed.T3 in our previous cell culture experiments without the CAAX tag [34, 43]. Here, we were able to verify the toxic effect of DsRed-CAAX in transgenic *D. suzukii* flies and while the mechanism is not solved, the CAAX tag seems to play a role perhaps by guiding the DsRed protein to the membrane and forming aggregates that disrupt the cell walls. This phenomenon could be used as a novel, universal effector tool for insect control programs. For example, when combined with a Tet-Off system for conditional expression, DsRed-CAAX could be deployed in a TESS to induce female-specific lethality. With such a universal effector gene in hand, SIT programs would not require the isolation of endogenous pro-apoptotic genes from every pest species. DsRed-CAAX could also be used as an internal marker to screen for potential loss of function mutations during mass rearing. This universal effector would also reduce the effort required to test transgenes for potential detrimental effects on human health or the environment. Furthermore, DsRed is an excellent candidate because it is widely used as a marker in plants, vertebrates, and invertebrates [52–57] but is not known to be toxic following oral administration, as shown by the analysis of predators feeding on mosquito larvae expressing a *DsRed* transgene [58]. It is also stable under laboratory and field conditions, making it a reliable candidate marker for SIT programs [59, 60].

Finally, 2A peptides could be used to develop effective Cas9-mediated gene drives to achieve efficient pest management in a wide variety of species. Such systems would rely on Cas9 cassettes that achieve drive via multi-generational homology-directed repair (HDR) together with marker genes for identification of the knock-in mutants [61]. For example, DsRed and GFP have been used as reporters to evaluate the efficiency of gene drive in mosquitos [62, 63]. For pest control, the male determination gene (*M*) was proposed as the effector gene to achieve population suppression in mosquitos [46], and resistance-suppressing effectors were proposed to achieve population replacement in the diamondback moth *Plutella xylostella* [64]. However, both the marker and effector genes would need additional regulatory elements, adding to the overall size of the gene-drive cassette, and the knock-in efficiency of Cas9 was inversely related to the insert size [26, 65, 66]. It is possible to co-express Cas9 with the marker and effector genes using 2A peptides, and the smaller insert size could therefore increase the efficiency of gene drive based on HDR. Indeed, a promoter-Cas9-2A-GFP cassette was used to generate transgenic mice and fruit flies that were not only fertile and able to breed to homozygosity, but also exhibited fluorescence and high CRISPR editing efficiency [67, 68]. Here we observed differences between the DrosCV-2A and TaV-2A peptides, and previous reports have also provided evidence that the sequence [26, 69] and position [17, 32] of proteins relative to the 2A peptide could influence the cleavage efficiency. The performance of a specific 2A peptide should therefore be tested for each construct, cell type, and organism, and the efficient synthesis of every effector protein in a multicistronic construct should be carefully monitored for different pest control strategies.

## Author contributions

JS, YY, and ZF performed research. JS, YY, ZF, and MFS designed experiments, analyzed data and wrote the paper.

## Acknowledgements

We thank Bashir Hosseini for excellent technical assistance. This work was supported by the Fraunhofer Attract program (‘Applications for population control of *D. suzukii’*; to MFS), the Emmy Noether program of the German Research Foundation (SCHE 1833/1-1; to MFS) and the LOEWE Center for Insect Biotechnology and Bioresources of the HMWK.

## Availability of data and materials

All data discussed in the paper will be made available to readers.

---

**Figure S1.**
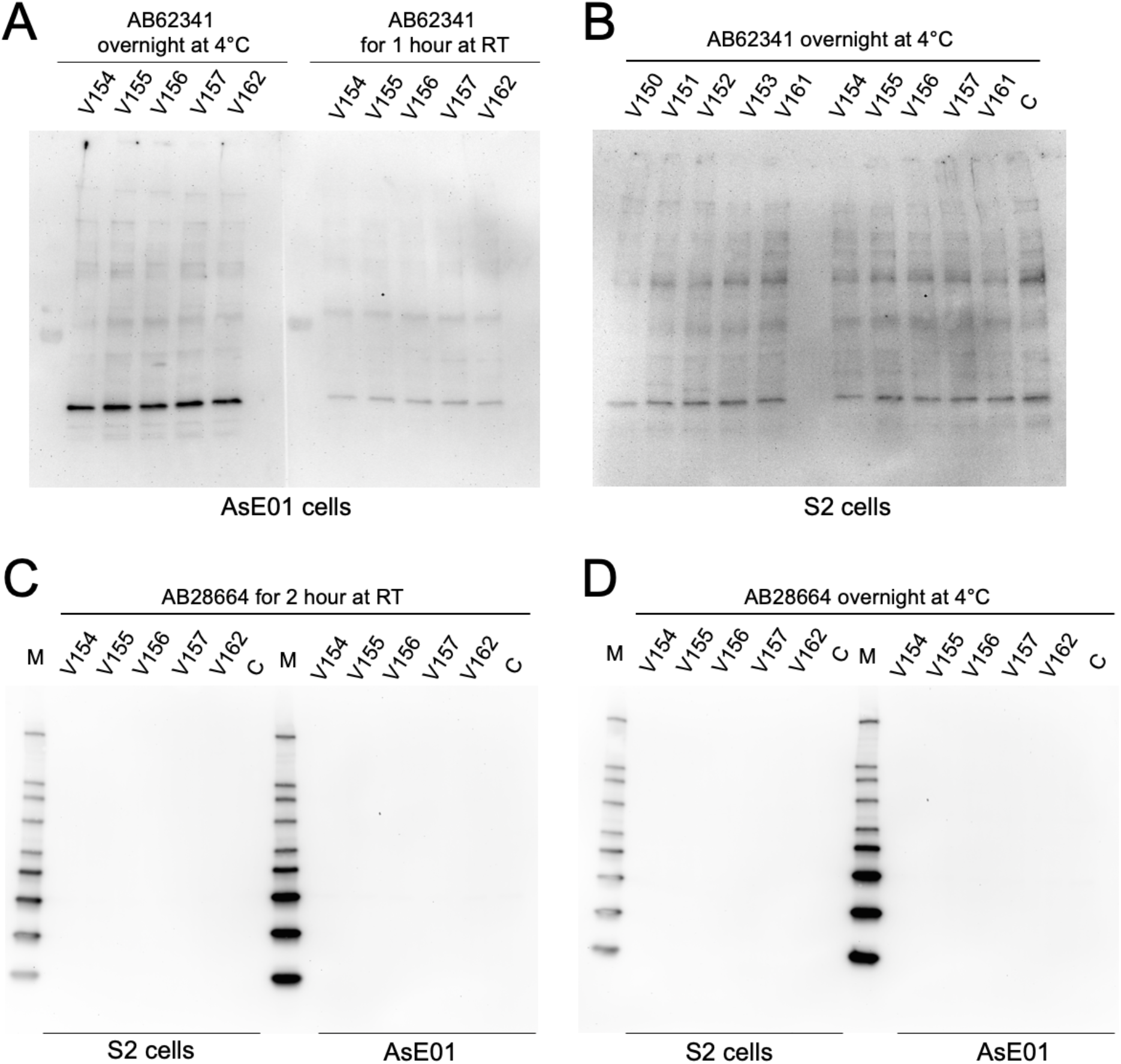
Additional western blot analysis to identify DsRed derivatives produced by cells transfected with pIE-4 constructs. (A). The DsRed-specific antibody AB62341 (ABCAM) was used to stain AsE01 cells at either 4°C overnight or room temperature (RT) for 1 h. Non-specific bands were observed for all constructs including the control construct (V162) without the 2A peptide, and the 4°C overnight incubation produced clearer results. (B). AB62341 was also used to stain S2 cells at 4°C overnight, and once again no specific DsRed products were detected. C = control, non-transfected cells. An alternative DsRed-specific antibody AB28664 (ABCAM) was used to stain both cell types at RT for 2 h (C) or 4°C overnight (D). No bands were detected. M = size markers.

**Figure S2.**
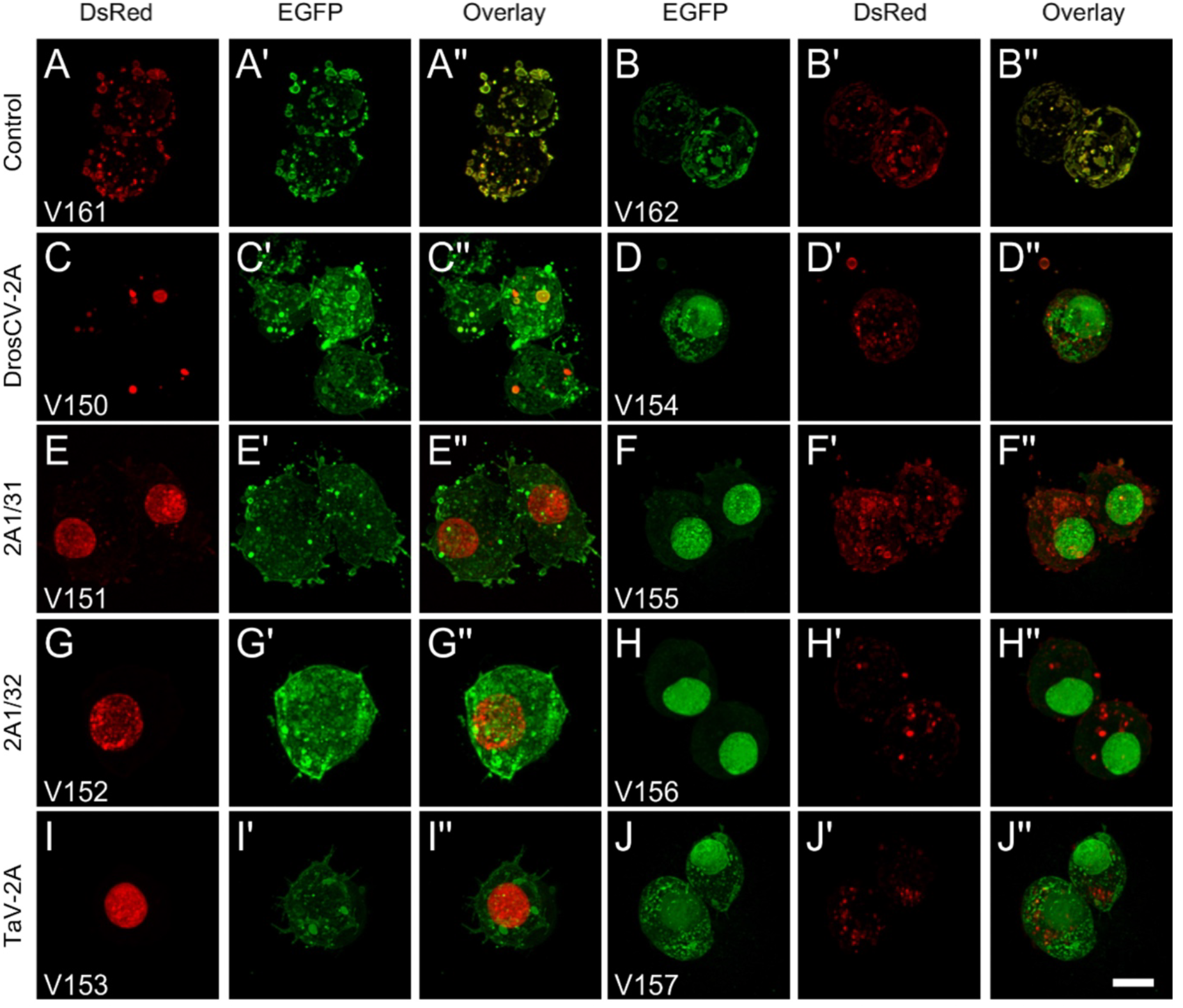
Expression of constructs containing a 2A peptide in *Drosophila* S2 cells. The control constructs V161 and V162 express a DsRed-EGFP fusion protein and hence show co-localization of the two proteins in an irregular pattern corresponding to the cell membrane and putative nuclear membrane. The constructs containing a 2A peptide (V150–V157, C-J′′) show independent localization of the two fluorescent proteins. Some of those constructs display additional effects. In the DrosCV-2A construct V150, DsRed-NLS was not homogenously distributed in the nucleus but accumulated in small vesicular structures with strong fluorescence. Vesicular structures around the cell membrane were also observed in the EGFP channel (C-C′′). In contrast to EGFP-CAAX, DsRed-CAAX (V154–V157) showed a punctate localization pattern in the cell membrane. Scale bar = 5 µm.

**Table S1.** Germ-line transformation of *D. suzukii*

| Vector (size) | Injected eggs | Hatched larvae | Fertile adults | Helper | Transgenic lines<sup>d</sup> | Transformation frequency | Hatch rate | Fertile eclosion rate (to larvae)<sup>e</sup> |
| --- | --- | --- | --- | --- | --- | --- | --- | --- |
| V220<br>(11,249 bp) | 665 | 85 | 29 | AH286<sup>a</sup> | 0 | 0 | 12.8% | 34.1% |
|  | 322 | 54 | 16 | AH286 | 0 | 0 | 16.8% | 29.6 % |
|  | 174 | 75 | 11 | AH286 | 0 | 0 | 43.1% | 14.7% |
|  | 440 | 17 | 9 | AH286 | 1 | 11.1% | 3.9% | 52.9% |
| V221<br>(11,252 bp) | 443 | 102 | 9 | AH286 | 0 | 0 | 23.0% | 8.8% |
|  | 728 | 137 | 14 | AH286/<br>V315<sup>b</sup> | 0 | 0 | 18.8% | 10.2% |
|  | 729 | 134 | 28 | AH286/<br>V315 | 1 | 3.6% | 18.4% | 20.9% |
| V222<br>(11,246 bp) | 571 | 163 | 15 | AH286/<br>V315 | 0 | 0 | 28.5% | 9.2% |
|  | 578 | 86 | 8 | AH286/<br>V315 | 0 | 0 | 14.9% | 9.3% |
|  | 608 | 163 | 125 | RNA-helper<sup>c</sup> | 0 | 0 | 26.8% | 76.7% |
|  | 749 | 152 | 11 | AH286/<br>V315 | 0 | 0 | 20.3% | 7.2% |
| V223<br>(11,249 bp) | 488 | 82 | 30 | AH286/<br>V315 | 2 | 6.7% | 16.8% | 36.6% |

<sup>a</sup> AH286 is the *phsp-pBac* transposase helper plasmid (Schetelig et al., 2013).
<sup>b</sup> V315 a vector containing a hyperactive transposase (Eckermann et al., 2018).
<sup>c</sup> *piggyBac* RNA helper was *in vitro* synthesized using AH286 as a template.
<sup>d</sup> Transgenic lines that derived from independent G<sub>0</sub> crosses.
<sup>e</sup> Number of fertile adults divided by number of hatched larvae.

**Table S2.**
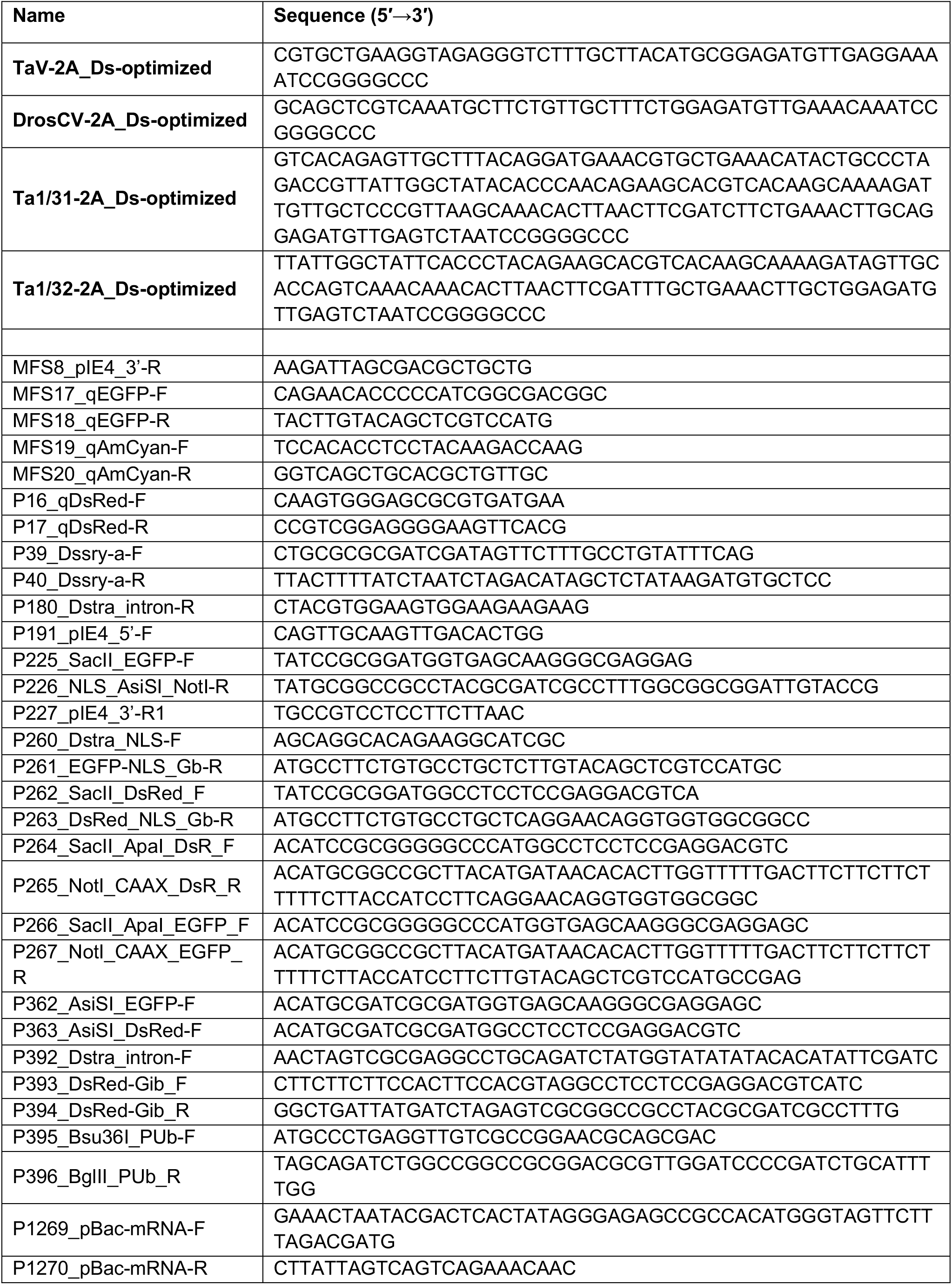
2A peptide and primer sequences

